# Effects of sex and estrous cycle on intravenous oxycodone self-administration and the reinstatement of oxycodone-seeking behavior in rats

**DOI:** 10.1101/2023.06.02.543393

**Authors:** Nicole M. Hinds, Ireneusz D. Wojtas, Corinne A. Gallagher, Claire M. Corbett, Daniel F. Manvich

**Affiliations:** Graduate School of Biomedical Sciences, Rowan University School of Osteopathic Medicine, Stratford, NJ, USA; Department of Cell Biology and Neuroscience, Rowan University School of Osteopathic Medicine, Stratford, NJ, USA

**Keywords:** oxycodone, opioids, self-administration, reinstatement, sex, estrous

## Abstract

The increasing misuse of both prescription and illicit opioids has culminated in a national healthcare crisis in the United States. Oxycodone is among the most widely prescribed and misused opioid pain relievers and has been associated with a high risk for transition to compulsive opioid use. Here, we sought to examine potential sex differences and estrous cycle-dependent effects on the reinforcing efficacy of oxycodone, as well as on stress-induced or cue-induced oxycodone-seeking behavior, using intravenous (IV) oxycodone self-administration and reinstatement procedures. In experiment 1, adult male and female Long-Evans rats were trained to self-administer 0.03 mg/kg/inf oxycodone according to a fixed-ratio 1 schedule of reinforcement in daily 2-hr sessions, and a dose-response function was subsequently determined (0.003-0.03 mg/kg/inf). In experiment 2, a separate group of adult male and female Long-Evans rats were trained to self-administer 0.03 mg/kg/inf oxycodone for 8 sessions, followed by 0.01 mg/kg/inf oxycodone for 10 sessions. Responding was then extinguished, followed by sequential footshock-induced and cue-induced reinstatement tests. In the dose-response experiment, oxycodone produced a typical inverted U-shape function with 0.01 mg/kg/inf representing the maximally effective dose in both sexes. No sex differences were detected in the reinforcing efficacy of oxycodone. In the second experiment, the reinforcing effects of 0.01-0.03 mg//kg/inf oxycodone were significantly attenuated in females during proestrus/estrus as compared to metestrus/diestrus phases of the estrous cycle. Neither males nor females displayed significant footshock-induced reinstatement of oxycodone seeking, but both sexes exhibited significant cue-induced reinstatement of oxycodone seeking at magnitudes that did not differ either by sex or by estrous cycle phase. These results confirm and extend previous work suggesting that sex does not robustly influence the primary reinforcing effects of oxycodone nor the reinstatement of oxycodone-seeking behavior. However, our findings reveal for the first time that the reinforcing efficacy of IV oxycodone varies across the estrous cycle in female rats.

## 1. Introduction

The misuse of prescription and illicit opioids has culminated in a national healthcare crisis in the United States (Baker et al., 2021; Volkow and Blanco, 2021; Volkow et al., 2022). Fatal overdoses involving opioids have increased more than threefold over the past decade (Centers for Disease Control and Prevention, 2021), with 77,766 deaths reported during the 12-month period ending December 2021, the highest ever recorded to date (Centers for Disease Control and Prevention, 2022). Prescription opioid pain relievers accounted for the overwhelming majority of opioids misused in 2020 (more than 90%) (Centers for Disease Control and Prevention, 2021), corroborating evidence that opioid use disorder (OUD) is commonly preceded either by prescribed treatment with opioid pain relievers or by the diversion of these medications from close contacts (McHugh et al., 2015; Centers for Disease Control and Prevention, 2021). In response to the ongoing opioid epidemic, research efforts aimed at understanding the neurobiological factors that confer enhanced susceptibility to opioid misuse have intensified, with the goal of identifying novel behavioral and/or pharmacological strategies to treat OUD or to prevent its development during prescribed opioid use (Volkow and Blanco, 2021).

Among the variables well-established to play a role in the vulnerability to opioid misuse is biological sex. Women are more likely than men to be prescribed opioid analgesics for the treatment of pain (Hirschtritt et al., 2018; Koons et al., 2018; Mazure and Fiellin, 2018), to self-report opioid misuse as a coping strategy to mitigate stress or negative affect (McHugh et al., 2013), and to experience more intense opioid craving (Back et al., 2011). Such factors are likely to partially mediate the accelerated progression to opioid misuse and OUD that has been reported in women (Anglin et al., 1987; Hser et al., 1987; Hernandez-Avila et al., 2004; Sartor et al., 2014). Moreover, these sex differences may be contributing to an exacerbated impact of the opioid epidemic on women, as rates of fatal opioid-involved overdoses since 1999 have accelerated faster in women than men (14-fold vs, 10-fold) (Centers for Disease Control and Prevention, 2021).

In agreement with clinical reports of sex differences in the abuse-related effects of opioids, several preclinical studies employing models of drug self-administration have reported enhanced opioid reinforcing efficacy in female subjects. This pattern has reliably been observed using various schedules of reinforcement (e.g., fixed-ratio, progressive-ratio), durations of daily access (e.g., short access, long access), and different mu opioid receptor agonists (heroin, fentanyl, morphine, remifentanil). In general, female subjects acquire opioid self-administration faster (Lynch and Carroll, 1999; Carroll et al., 2002; Cicero et al., 2003; Thorpe et al., 2020; Malone et al., 2021), exert greater effort for opioid reinforcement or opioid seeking (Cicero et al., 2003; Bakhti-Suroosh et al., 2021; Towers et al., 2022), and self-administer more opioid reinforcers than males, especially at low unit doses that typically encompass the ascending limb of the inverted U-shaped dose-response function (Alexander et al., 1978; Klein et al., 1997; Cicero et al., 2003; Lacy et al., 2016; Towers et al., 2019; Townsend et al., 2019; Bakhti-Suroosh et al., 2021; Towers et al., 2022; but see Stewart et al., 1996).

Oxycodone is among the most widely prescribed and misused opioid pain relievers (Centers for Disease Control and Prevention, 2021) and has been associated with a high risk for transition to compulsive opioid misuse (Rosenblum et al., 2007; Remillard et al., 2019). Consequently, oxycodone has been implicated as a major contributor to the current opioid epidemic (Compton and Volkow, 2006; Hedegaard et al., 2018; Kibaly et al., 2021; Rivas et al., 2021). Yet despite oxycodone’s prominent role in the opioid crisis and the aforementioned influence of biological sex on opioid effects, sex differences in oxycodone reinforcement specifically have only recently been explored in a small number of preclinical studies, with equivocal results. For example, studies using oral oxycodone self-administration procedures in rats and mice have demonstrated higher oxycodone intake in female subjects (Fulenwider et al., 2020; Phillips et al., 2020; Zanni et al., 2020), and greater IV oxycodone reinforcement has been observed in female rats at high doses of an ascending dose-response determination (Mavrikaki et al., 2017). Moreover, escalation of extended access IV oxycodone self-administration occurred more rapidly and to greater effect in female vs. male rats (Kimbrough et al., 2020). By contrast, other studies have failed to detect sex differences in IV oxycodone self-administration when a single dose was held constant across sessions under short access conditions (1-4 h/session) (Mavrikaki et al., 2017; Collins et al., 2020; Mavrikaki et al., 2021), and enhanced oxycodone reinforcement has been reported in males as compared to females under conditions of food restriction (Mavrikaki et al., 2017). Direct comparisons between these experiments are difficult due to parametric differences between them including but not limited to species, strains, schedules of reinforcement, session durations, oxycodone doses/concentrations, and housing/feeding conditions, each of which may have contributed variability to the experimental results. Further complicating this narrative is recent evidence that fluctuations in ovarian hormones across the estrous cycle modulate the reinforcing efficacy of heroin in female rats (Lacy et al., 2016; Smith et al., 2021; Smith et al., 2022) (but see Stewart et al., 1996; Roth et al., 2002). However, the impact of estrous cycle phase on oxycodone self-administration has received little attention to date, with only a single study reporting no effect of estrous cycle on oral oxycodone self-administration (Fulenwider et al., 2020). Importantly, the impact of estrous cycle on IV oxycodone reinforcement or oxycodone seeking following IV self-administration has not yet been explored.

The objective of the present study was to further characterize the roles of sex and estrous cycle in the abuse-related effects of oxycodone using preclinical rodent models of IV drug intake and relapse. In the first experiment, we assessed whether there are sex differences in the primary reinforcing effects of IV oxycodone. Male and female rats were initially trained to self-administer 0.03 mg/kg/inf oxycodone under a fixed-ratio 1 (FR1) schedule of reinforcement, and then dose-response functions were determined (0.003 – 0.03 mg/kg/inf) and compared between sexes. In the second experiment, male and female rats were allowed to self-administer 0.03 mg/kg/inf for 8 sessions followed by access to 0.01 mg/kg/inf for 10 sessions, and reinforcing efficacy of oxycodone was compared between sexes and across estrous cycle phases in females. We also assessed in these subjects whether the magnitude of stress- or cue-induced reinstatement of extinguished oxycodone-seeking behavior varied as a function of either sex or estrous cycle.

## 2. Materials and Methods

### 2.1 Animals

Subjects were male (200-225 g upon arrival) and female (175-200 g upon arrival) adult Long-Evans rats acquired from Charles River Laboratories (Wilmington, MA) or Envigo (Indianapolis, IN). Animals were housed in polycarbonate cages (43 x 24 x 20 cm) under a reverse 12-h light-dark cycle (lights off at 09:00am) prepared with one wooden chew block per animal and *ad libitum* access to rodent chow and water. Rats were initially pair-housed upon arrival in same-sex pairs and were allowed to acclimate to the facility for one week prior to onset of experiments. All procedures were conducted in accordance with the NIH Guide for the Care and Use of Laboratory Animals and were approved by the Rowan University Institutional Animal Care and Use Committee. Behavioral studies took place during the dark phase of the light cycle (between 10:00am – 04:00pm).

### 2.2 Food training and surgery

Following acclimation to the facility, animals were trained to lever-press for 45 mg food pellets (Bio-Serv, Flemington, NJ) in operant chambers housed within ventilated, sound-attenuating cubicles (Med Associates Inc., St Albans, VT). Each training session began with the extension of two levers (right, left) into the operant chamber. A response on the active lever (randomly assigned and counterbalanced across subjects) delivered a single food pellet into a receptacle located in the center of the operant panel, equidistant between the two levers. Food training sessions were terminated if an animal earned 100 reinforcers or 4 h elapsed, whichever occurred first. Animals were deemed trained once they earned ≥ 60 reinforcers and emitted ≥ 70% of total responses on the active lever within a single session, typically achieved within 1-3 sessions. If a rat failed to satisfy these criteria within 4 sessions, food training was halted and the animal progressed on to surgical catheter implantation.

Following food training sessions, animals were prepared with chronic indwelling IV catheters as described previously (Schroeder et al., 2010). Briefly, under inhaled isoflurane anesthesia, a custom-fabricated silastic catheter (P1 Technologies, Roanoke, VA) was inserted into the right jugular vein and secured to the vessel using nonabsorbable suture. The catheter passed subcutaneously to an infusion port positioned between the scapulae from which an infusion cannula extended through the skin. Beginning on the day of surgery and occurring regularly thereafter, catheters were flushed 5-6 days per week with 0.05 ml of gentamicin (4 mg/ml) and locked with 0.1 ml of heparin (300 units/ml). When not in use, catheters were protected with a silastic obturator and a stainless-steel dust cap. Animals were singly-housed following surgery and for the remainder of the study and were allowed a minimum of one week of recovery prior to the onset of oxycodone self-administration experiments.

### 2.3 Intravenous oxycodone self-administration

Oxycodone self-administration was conducted 5-7 days/week in 2 h sessions under a FR1 schedule of reinforcement. Each session began with the extension of the two levers into the operant chamber and illumination of a house light located on the opposite wall which served as a discriminative stimulus to indicate reinforcer availability. An automated syringe pump (PHM100; Med Associates) was located outside the cubicle and connected via Tygon tubing to a 22-g liquid swivel (Instech, Plymouth Meeting, PA) that was secured by a counterweighted arm above the operant chamber. A metal tether protected an additional segment of Tygon tubing that extended from the output of the liquid swivel to the vascular access port of the subject in the operant chamber. Med-PC V software (Med Associates) interfaced with each operant chamber to control all outputs and record lever presses. Subject weights were entered into the self-administration program daily and were used to calculate pump durations to ensure accuracy of oxycodone unit doses delivered to each animal.

During self-administration sessions, a response on the active lever resulted in an infusion of oxycodone that was paired with illumination of the cue light above the active lever and termination of the house light for 20 s (timeout), during which active lever responses were recorded but had no scheduled consequences. Inactive lever presses throughout the session were also recorded but had no consequences. Animals were provided a single wooden chew block during all operant-behavioral sessions to mitigate gnawing of operant chamber components, a behavioral output which we observed in pilot studies of oxycodone self-administration and which has been reported by others (Zanni et al., 2020). Sessions were terminated if animals earned 60 infusions or if 2 h elapsed, whichever occurred first. If animals earned 60 infusions, the levers were retracted and cue/house lights were turned off, but the animal remained in the operant chamber for the remainder of the 2 h session duration.

### 2.4 Experiment 1: Oxycodone dose-response function – role of sex

12 male and 10 female rats served as subjects for Experiment 1. Following food training and surgery, rats were initially allowed to self-administer 0.03 mg/kg/inf oxycodone in daily sessions. 0.03 mg/kg/inf was selected as the first dose of the dose-response determination based on pilot studies in our laboratory showing that rats reliably acquired self-administration at this dose, defined as ≥ 15 reinforcers earned across 3 consecutive sessions. Responding for 0.03 mg/kg/inf was deemed stable when the following two criteria were met across four consecutive sessions: < 30% variability in active lever response rate, and < 20% variability in the number of reinforcers earned. A dose-response was subsequently determined using a descending dose-order procedure, such that animals were next tested at 0.01 mg/kg/inf followed by 0.003 mg/kg/inf. In order to transition from one dose to the next, the aforementioned stabilization criteria had to be satisfied. Once all three doses had been assessed, rats underwent extinction training during which active lever presses resulted in saline infusions without presentation of the oxycodone-paired cue light. Saline-maintained self-administration was considered stable if a rat met the stabilization criteria described above, or if they emitted ≤ 20 active lever presses across two consecutive sessions. 2 male and 3 female rats were ultimately excluded from the final analysis due to failure to acquire oxycodone self-administration (1 male, 1 female) or loss of catheter patency during the experiment (1 male, 2 females). Therefore, the final cohort consisted of 10 male and 7 female subjects.

### 2.5 Experiment 2: Oxycodone self-administration, extinction, and reinstatement – role of sex and estrous cycle

10 male and 23 female rats, distinct from those used in Experiment 1, initially served as subjects for Experiment 2. Following food training and surgery, these rats were allowed to self-administer 0.03 mg/kg/inf oxycodone for 8 sessions under the FR1 schedule of reinforcement. The unit dose of oxycodone was then reduced to 0.01 mg/kg/inf and animals self-administered this dose for a further 10 sessions. 2 male and 5 female rats were removed from the study due to loss of catheter patency prior to conclusion of the self-administration portion of the experiment, and 1 additional female rat was removed because it failed to acquire oxycodone self-administration. Therefore, a total of 8 male and 17 female rats completed the self-administration segment of Experiment 2.

Beginning on session 19, responding was extinguished in 2 h sessions (once per day, 5-7 days/week) during which the house light was illuminated but neither active nor inactive lever presses had scheduled consequences. For each individual rat, responding was deemed extinguished when its rate of responding on the active lever in each of two consecutive extinction sessions reached < 30% of the mean active lever response rate during its final 3 sessions of 0.01 mg/kg/inf oxycodone self-administration. Animals that failed to meet this extinction criterion within 21 sessions were excluded from reinstatement experiments (2 males, 1 female). Therefore, the final cohort that progressed to reinstatement testing consisted of 6 male and 16 female subjects. In some cases, extinction training was extended in females by 1-2 sessions beyond the satisfaction of extinction criteria in order to ensure testing during a specific phase of the estrous cycle, so as to maintain equivalent sample sizes between experimental groups. Subjects were still required to meet extinction criteria in order to undergo a reinstatement test. On days where a specific estrous cycle phase was targeted, experimenters used prior days’ swabs to estimate upcoming cycle phases and verified that the subject was in the targeted phase on the morning of the reinstatement test session.

Animals underwent a stress-induced reinstatement test on the day immediately following satisfaction of extinction criteria (females also had to be in their targeted estrous cycle phase). Rats were placed in an operant chamber, with levers retracted and all lights off, and exposed to 15 min of intermittent, unpredictable footshock (0.5 mA, 0.5 s duration, 3 – 80 s variable intervals). These parameters for footshock exposure were chosen because they reliably induce reinstatement of drug seeking for several drugs of abuse (Shaham et al., 1997; Schank et al., 2011; Schroeder et al., 2013; Ogbonmwan et al., 2015; Fulenwider et al., 2020). Three min after the final footshock was delivered, levers were extended and the house light was illuminated, and responses were recorded under extinction conditions for 2 h. After footshock-induced reinstatement testing, animals were given additional extinction sessions until aforementioned extinction criteria were again satisfied. On the day following re-establishment of extinction, animals underwent a 2 h cue-induced reinstatement test during which active lever presses resulted in the illumination of the oxycodone-paired 20 s cue light located above the active lever.

Early results obtained during footshock-induced reinstatement tests and cue-induced reinstatement tests suggested that levels of operant responding were not above extinction levels in a small number of subjects. To rule out the possibility that exposure to footshock within the self-administration context produced a nonspecific suppression of operant behavior, we sought to determine whether animals would resume lever-pressing under drug-reinforced conditions. Following the conclusion of all reinstatement tests, a subset of animals that had maintained catheter patency through the conclusion of reinstatement testing (6 males, 10 females) underwent a final round of extinction sessions after the cue-induced reinstatement test and were allowed to self-administer 0.01 mg/kg/inf oxycodone in a single “reacquisition” session using parameters identical to earlier self-administration sessions.

### 2.6 Estrous Cycle Monitoring

Estrous cycle was monitored in female rats that served as subjects in Experiment 2 (n=17). Animals were habituated to vaginal lavaging for 3-4 days prior to self-administration onset, with sample collection beginning on the first day of self-administration. Vaginal swabs were performed 1-2 hours prior to each operant-behavioral session to mitigate any impact of handling/swabbing on operant-behavioral output while also attempting to minimize phase transition during the interval between sample collection and session onset. Samples were collected using procedures adapted from those reported previously (Bangasser and Shors, 2008; Corbett et al., 2021). Briefly, while gently restraining the rat, a cotton-tipped applicator was rapidly dipped in sterile saline, blotted until slightly damp, and then inserted into the vaginal canal approximately 0.5 cm and rotated for 4-5 seconds. The applicator was then pressed onto a glass microscope slide using a rolling motion and allowed to air dry. Slides were subsequently stained with 1% toluidine blue and examined by light microscopy at 10x-20x by a trained experimenter who was blinded to the animals’ treatment conditions. Estrous cycle stages were identified based on the presence and morphology of cells as described previously (Hubscher et al., 2005; Goldman et al., 2007; Cora et al., 2015). Each sample was classified as one of the following four stages: metestrus, classified by approximately similar amounts of nucleated epithelial cells, anucleated cornified epithelial cells, and leukocytes; diestrus, classified based on predominance of leukocytes with occasional epithelial cells; proestrus, classified by ≥75% of nucleated epithelial cells; estrus, classified by ≥75% of anucleated cornified epithelial cells. If a sample was collected during a phase transition in which none of the above criteria were satisfied, the sample was assigned to the later of the two phases that were apparent in the sample. At the conclusion of Experiment 2, three of sixteen female rats were removed from final data analyses because their vaginal cytology was irregular and lavages did not produce cells in sufficient quantities to allow for rigorous estrous cycle phase determinations.

### 2.7 Drugs

Oxycodone hydrochloride was provided by the National Institute on Drug Abuse Drug Supply Program (Research Triangle Park, NC) and dissolved at concentrations of 0.018 – 0.18 mg/ml in sterile bacteriostatic saline, passed through a 0.22 μm filter, and stored in sterile vials at 4°C. All doses are reported as the salt weight.

### 2.8 Statistical analyses

Experiment 1: The number of sessions to reach stabilization, active and inactive lever presses, infusions earned, and rates of responding on active and inactive levers were analyzed using two-way mixed-factors ANOVA with repeated measures on dose and independent measures on sex. Post hoc multiple comparisons were performed using Dunnett’s tests or Tukey’s tests as specified. Number of sessions to reach food-training criteria were analyzed via unpaired t-test.

Experiment 2a – estrous cycle and oxycodone self-administration: Active lever presses, inactive lever presses, infusions earned, and rates of responding on active and inactive levers were analyzed using two-way repeated measures ANOVA with repeated measures on both dose and estrous cycle. Because each factor consisted of only two levels, post hoc analyses following significant main effects were not required. Number of sessions to reach food-training criteria were analyzed via unpaired t-test. Body mass was analyzed independently in each sex using a one-way repeated measures ANOVA with session as the within-subjects factor.

Experiment 2b – footshock-induced reinstatement, cue-induced reinstatement, and reacquisition of oxycodone self-administration: Active and inactive lever presses were each analyzed using two-way mixed-factors ANOVA with repeated measures on experimental phase (extinction vs. reinstatement, or extinction vs. reacquisition) and independent measures on group. Because the only significant effects identified in these analyses were main effects of experimental phase for which there are only two levels, post hoc analyses were not required. Number of sessions to reach extinction criteria were analyzed via one-way ANOVA or unpaired t-test as specified with group as a between-subjects factor.

Data were plotted and analyzed using GraphPad Prism version 9.2 (GraphPad Software, La Jolla, CA, USA). Significance was set at α = 0.05 for all statistical tests.

## 3. Results

### 3.1 IV oxycodone is equally effective as a reinforcer in male and female rats under FR1

10/10 male rats (100%) and 6/7female rats (~86%) used in Experiment 1 achieved lever-press training under food reinforcement within 1-3 sessions with no significant sex difference in the number of sessions to criteria (t_(14)_ = 0.17, p = 0.865) (**Fig. S1A**). The reinforcing effects of 0.003-0.03 mg/kg/inf IV oxycodone were then assessed in male and female rats under a FR1 schedule of reinforcement. Following acquisition and stabilization of 0.03 mg/kg/inf oxycodone self-administration, animals were subsequently allowed to self-administer oxycodone at 0.01 and 0.003 mg/kg/inf, presented in descending order, followed finally by saline substitution. Animals reached stabilization criteria within 4-14 sessions for each unit dose of oxycodone and 4-19 sessions for saline substitution (**Fig. 1A**). The number of sessions required to reach stable responding did not differ by dose (F_(3,45)_ = 0.43, p = 0.733) or sex (F_(1,15)_ = 0.11, p = 0.748), nor was there a significant dose × sex interaction (F_(3,45)_ = 0.93, p = 0.436). IV oxycodone self-administration resulted in an inverted U-shaped dose-response function, with 0.01 mg/kg/inf representing the maximally effective dose in both male and female rats (**Fig. 1B**). Statistical analysis of active lever presses revealed a significant main effect of oxycodone dose (F_(3,45)_ = 18.42, p < 0.0001), with no significant effect of sex (F_(1,15)_ = 2.77, p = 0.117) nor a dose × sex interaction (F_(3,21)_ = 1.97, p = 0.149). Post hoc Dunnett’s tests indicated that, collapsed across both sexes, responding on the active lever was significantly greater when any dose of oxycodone was available as compared to saline-maintained responding (p < 0.05). Rats also emitted higher inactive lever presses when 0.003 or 0.01 mg/kg/inf oxycodone was available as compared to saline (**Fig. 1B**) (main effect of dose, F_(3,451)_ = 3.97, p = 0.014; post hoc Dunnett’s tests, p < 0.05), however there was no effect of sex on inactive lever responding (F_(1,15)_ = 0.21, p = 0.653) nor was there a dose × sex interaction (F_(3,45)_ = 0.03, p = 0.992). Because a subset of subjects earned the maximum allowable 60 infusions when 0.003 or 0.01 mg/kg/inf oxycodone were available for self-administration, we further examined potential sex differences in oxycodone reinforcement by replotting the data using response rate (number of responses ÷ total session time) as the dependent measure of reinforcing efficacy, which adjusts for early termination of sessions due to maximum reinforcer delivery. This reanalysis also resulted in an inverted U-shaped dose-response function with a peak at 0.01 mg/kg/inf oxycodone (**Fig. 1C**). Statistical analysis of active lever response rate revealed a significant main effect of dose (F_(3,45)_ = 12.61, p < 0.0001) with no main effect of sex (F_(1,15)_ = 2.66, p = 0.124) nor a dose × sex interaction (F_(3,45)_ = 0.67, p = 0.577). Post hoc Dunnett’s tests indicated that, collapsed across both sexes, 0.003 and 0.01 mg/kg/inf oxycodone maintained higher rates of responding on the active lever than saline (p < 0.05). Analysis of response rate on the inactive lever revealed a main effect of oxycodone dose (F_(3,45)_ = 4.67, p = 0.006) but not of sex (F_(1,15)_ = 0.30, p = 0.591), nor was there a significant dose × sex interaction (F_(3,45)_ = 0.18, p = 0.908). Post hoc comparisons on the main effect of dose indicated that responding was higher on the inactive lever during self-administration of 0.003 or 0.01 mg/kg/inf oxycodone as compared to saline (p < 0.05). Finally, the number of oxycodone infusions varied as a function of dose (F_(2,30)_ = 6.94, p = 0.003) but there was not a significant main effect of sex (F_(1,15)_ = 0.32, p = 0.580) nor was there a significant dose × sex interaction (F_(2,30)_ = 0.32, p = 0.728) (**Fig. 1D**). Post hoc Tukey’s tests on the main effect of oxycodone dose indicated that animals self-administered less infusions when the highest dose of 0.03 mg/kg/inf oxycodone was available as compared to either the 0.003 or 0.01 mg/kg/inf doses. Taken together, these results suggest that IV oxycodone under a FR1 schedule of reinforcement engenders a typical inverted U-shaped dose-response function in both male and females rats without apparent sex differences in oxycodone efficacy.

**Figure 1:**
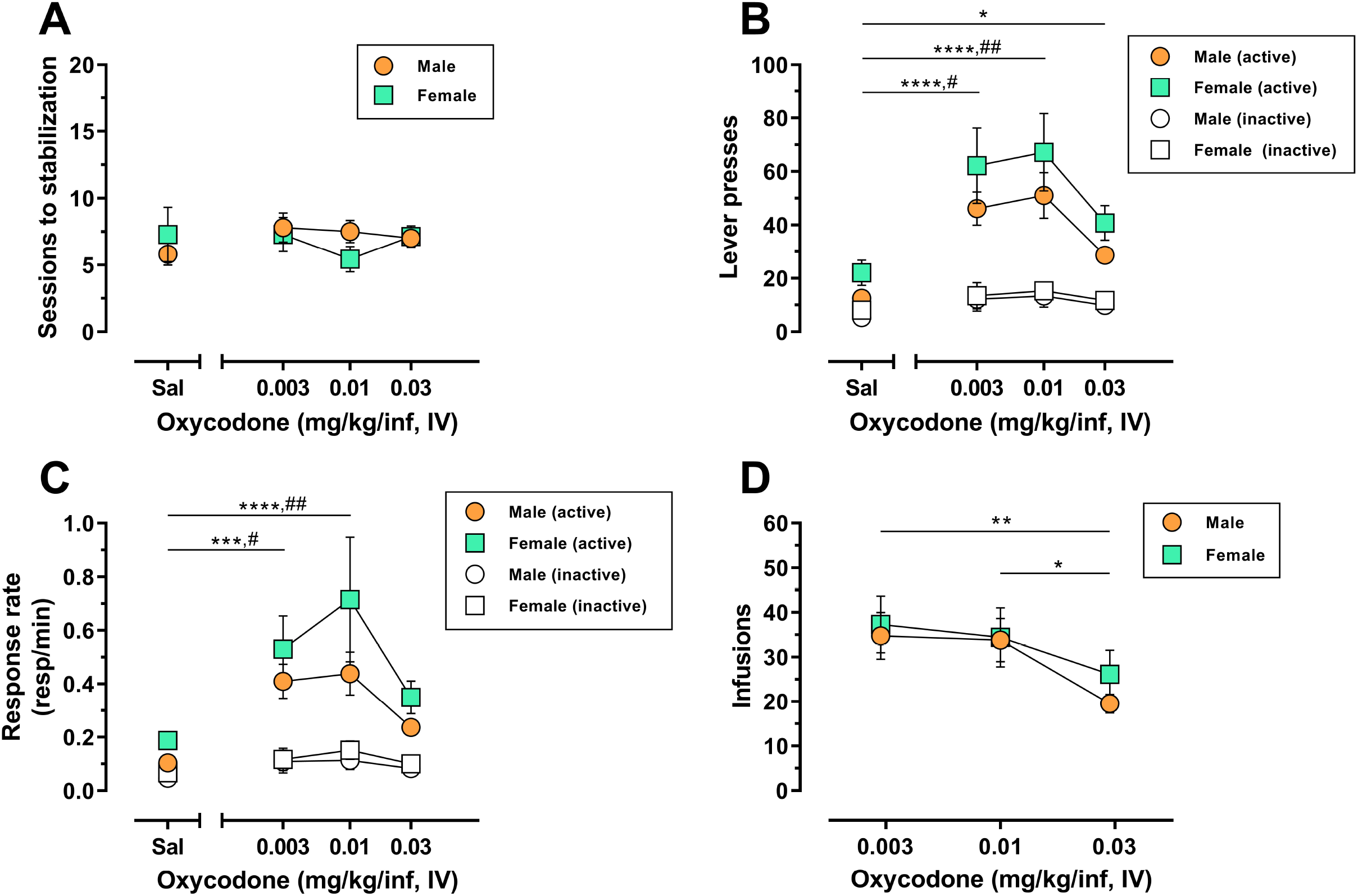
Dose-response determination for IV oxycodone self-administration in male and female rats. Male (n = 10) and female (n = 7) Long-Evans rats were trained to self-administer 0.03 mg/kg/inf oxycodone under a FR1 schedule of reinforcement followed by a dose-response determination. (**A**) Number of sessions required to reach stable self-administration at each unit dose of oxycodone. (**B**) Active lever presses (filled symbols) and inactive lever presses (empty symbols) maintained by each dose of oxycodone. (**C**) Rates of responding (responses per minute) on the active lever (filled symbols) and inactive lever (empty symbols) maintained by each dose of oxycodone. (**D**) Number of infusions earned at each dose of oxycodone. All data are presented as mean ± SEM values. * p < 0.05, ** p < 0.01, **** p < 0.0001, Dunnett’s test comparing active lever responding maintained by oxycodone vs. saline collapsed across sex (**B-C**) or Tukey’s test comparing oxycodone doses collapsed across sex (**D**). # p < 0.05, ## p < 0.01, Dunnett’s test comparing inactive lever responding maintained by oxycodone vs. saline collapsed across sex (**B-C**). “Sal”, saline. Absence of error bars indicates that SEM values did not extend beyond the limits of the depicted symbol.

### 3.2 Oxycodone-maintained responding is attenuated during proestrus/estrus in female rats during short-access FR1 self-administration

We next sought to determine whether oxycodone self-administration varies across the estrous cycle in female rats. Because our dose-response determination in Experiment 1 demonstrated that most rats acquire oxycodone self-administration at 0.03 mg/kg/inf and identified 0.01 mg/kg/inf oxycodone as the peak of the dose-response function in both sexes, we chose to assess the impact of estrous phase on the reinforcing efficacy of each of these oxycodone doses. Following lever-press training with food reinforcement and catheter implantation, male (n=8) and female (n=14) rats were trained on a FR1 schedule of reinforcement to self-administer 0.03 mg/kg/inf oxycodone for 8 sessions, followed by self-administration of 0.01 mg/kg/inf oxycodone for 10 sessions. Vaginal swabs were collected 1-2 h prior to each session in female subjects to determine their estrous cycle phase.

7/8 male rats (~88%) and 10/14 female rats (~71%) achieved lever-press training under food reinforcement within 1-3 sessions. There was no significant sex difference in the number of sessions to criteria for subjects that acquired (t_(15)_ = 1.93, p = 0.072) (**Fig. S1B**). Although female rats appeared to consistently exert higher levels of oxycodone-maintained active lever responding across both doses of oxycodone as compared to males (**Fig. 2A**), the main effect of sex on this measure was not significant, consistent with findings from Experiment 1 (main effect of sex, F_(1,20)_ = 1.09, p = 0.31; main effect of session, F_(17,340_ = 3.64, p < 0.0001; session × sex interaction, F_(17,340)_ = 0.64, p = 0.873). Similar to active lever responses, the number of oxycodone infusions earned varied by session (F_(17,340_ = 7.28, p < 0.0001) but did not differ by sex (F_(1,20_ = 0.47, p = 0.500) nor was there a session × sex interaction (F_(17,340_ = 0.46, p = 0.969) (**Fig. 2B**). Inactive lever responding did not differ by session (F_(17,340)_ = 1.25, p = 0.225) or sex (F_(1,20)_ = 0.11, p = 0.742), nor was there a session × sex interaction (F_(17,340)_ = 1.40, p = 0.133) (**Fig. 2A**). Over the duration of IV oxycodone self-administration exposure, body mass continuously and significantly increased in both male rats (F_(17,119)_ = 110.9, p < 0.0001) and female rats (F_(17,221)_ = 33.6, p < 0.0001), with males increasing body mass on average by 19.6% and females on average by 9.7% from session 1 to session 18 (**Fig. S2**).

**Figure 2:**
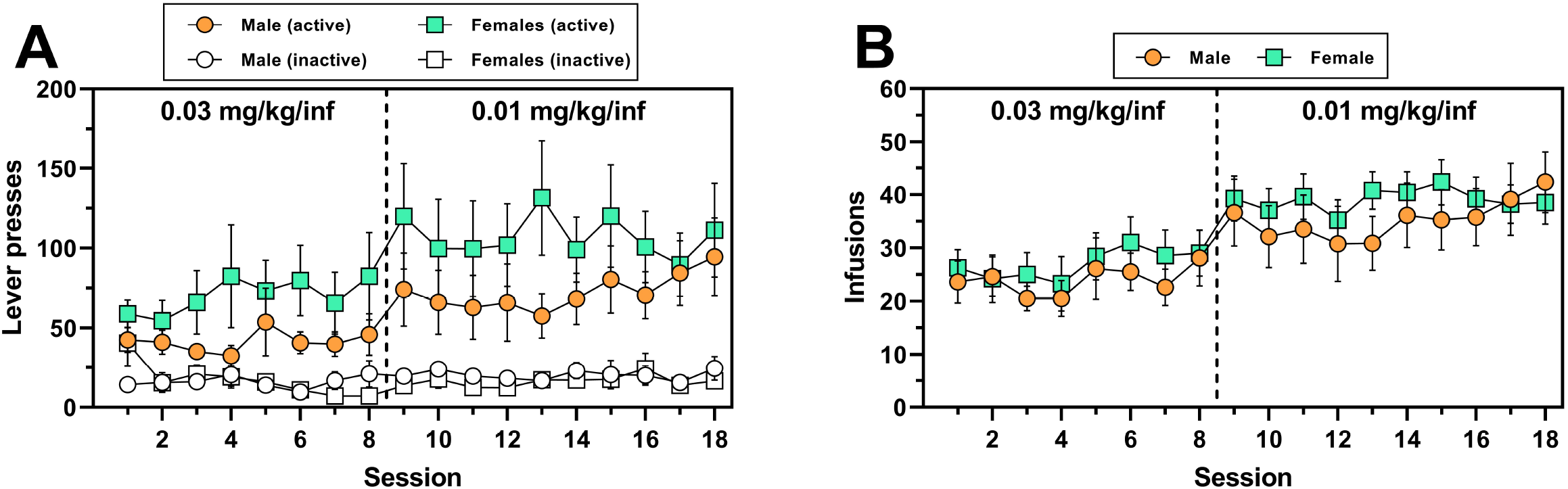
IV oxycodone self-administration behavior in male and female rats used in Experiment 2. Male (n = 8) and female (n = 14) Long-Evans rats self-administered 0.03 mg/kg/inf oxycodone for 8 sessions (sessions 1-8) followed by 10 sessions at 0.01 mg/kg/inf (sessions 9-18) under a FR1 schedule of reinforcement. (**A**) Active and inactive lever presses (filled and empty symbols, respectively) and (**B**) number of oxycodone infusions earned across sessions. The vertical dashed line in **A** and **B** indicates the transition from 0.03 to 0.01 mg/kg/inf oxycodone availability. Data are presented as mean ± SEM values.

The female rats included in the final analysis for Experiment 2 (n=14) displayed normal estrous cyclicity throughout the study without evidence of cycle dysregulation, similar to what has been reported previously following short-access IV oxycodone self-administration (Mavrikaki et al., 2017). We found in Experiment 1 that animals required 6-8 sessions on average to stabilize at 0.03 mg/kg/inf oxycodone self-administration and 6-10 sessions on average to stabilize at 0.01 mg/kg/inf oxycodone self-administration (**Fig. 1A**), with greatest variability during the first 4 sessions of self-administration at each dose. Because these initial sessions were characterized by unstable drug intake as animals learned the operant-behavioral contingencies and/or became accustomed to the pharmacological effects of the drug (or a change of unit dose), we chose to omit the first 4 sessions at each oxycodone unit dose from our assessment of estrous cycle impact on oxycodone reinforcement in Experiment 2, in order to reduce variability in the dataset and increase the rigor of the analysis. Therefore, with the first 4 sessions removed from each dose determination in Experiment 2, estrous cycle-dependent effects on oxycodone self-administration were examined in sessions 5-8 of 0.03 mg/kg/inf oxycodone availability and in sessions 5-10 of 0.01 mg/kg/inf oxycodone availability. We also excluded data from any sessions that immediately followed a day off from oxycodone self-administration due to several observed incidents of “rebound” response bursts that might have obfuscated the influence of estrous cycle on oxycodone reinforcement. The number of animals contributing self-administration data to each possible combination of estrous cycle phase and oxycodone dose are reported in **Table 1**. Animals were predominantly assessed during estrus or diestrus, with metestrus and proestrus detected at lower frequencies. Because an analysis comparing oxycodone reinforcement in each of the four cycle phases would have been underpowered as a result of the infrequent detection of proestrus and metestrus phases, we combined proestrus and estrus into one experimental dataset (P/E) and metestrus and diestrus into a second experimental dataset (M/D) in accordance with previous studies (Calipari et al., 2017; Brady et al., 2019; Johnson et al., 2019; Arguelles et al., 2021; Arguelles et al., 2022).

**Table 1.** Number of female rats assessed in proestrus, estrus, metestrus (diestrus I), or diestrus II during self-administration of 0.03 or 0.01 mg/kg/inf oxycodone in Experiment 2.

| Estrous Cycle Phase | # of Females Contributing Data (Percent Total) |
| --- | --- |
| *0.03 mg/kg/inf oxycodone* |  |
| Proestrus | 03/14 (21%) |
| Estrus | 14/14 (100%) |
| Metestrus/Diestrus I | 05/14 (36%) |
| Diestrus II | 14/14 (100%) |
| *0.01 mg/kg/inf oxycodone* |  |
| Proestrus | 03/14 (21%) |
| Estrus | 13/14 (93%) |
| Metestrus/Diestrus I | 05/14 (36%) |
| Diestrus II | 14/14 (100%) |

Two-way repeated-measures ANOVA for active lever presses (**Fig. 3A**) revealed that during IV oxycodone self-administration, females responded significantly less during proestrus/estrus as compared to metestrus/diestrus (main effect of estrous phase, F_(1,13)_ = 7.04, p = 0.020). There was also a significant main effect of dose on active lever responding (F_(1,13)_ = 8.32, p = 0.013) with no significant dose × estrous phase interaction detected (F_(1,13)_ = 1.59, p = 0.230). A similar pattern of results was obtained following the analysis of inactive lever responding (main effect of estrous phase, F_(1,13)_ = 6.49, p = 0.024; main effect of oxycodone dose, F_(1,13)_ = 16.57, p = 0.001; dose × estrous phase interaction, F_(1,13)_ = 1.00, p = 0.336) (**Fig. 3B**). As in Experiment 1, we reanalyzed oxycodone reinforcement using response rate as the dependent measure to correct for instances when lever-press availability was terminated early due to maximum reinforcer delivery. Two-way repeated measures ANOVA for active lever response rate revealed a significant main effect of both estrous phase (F_(1,13)_ = 13.19, p = 0.003) and oxycodone dose (F_(1,13)_ = 7.01, p = 0.020) with no significant dose × estrous phase interaction (F_(1,13)_ = 0.50, p = 0.493) (**Fig. 3C**). Similar results were obtained from the analysis of inactive lever response rate (main effect of estrous phase, F_(1,13)_ = 5.48, p = 0.036; main effect of oxycodone dose, F_(1,13)_ = 17.39, p = 0.001; dose × estrous phase interaction, F_(1,13)_ = 0.87, p = 0.368) (**Fig. 3D**). While oxycodone-maintained responding varied as a function of unit dose and estrous phase, the average number of oxycodone infusions earned did not significantly differ by oxycodone dose (F_(1,13)_ = 4.21, p = 0.061) or estrous phase (F_(1,13)_ = 0.30, p = 0.594) nor was there a dose × estrous phase interaction (F_(1,13)_ = 0.13, p = 0.728) (**Fig. 3E**). Collectively, these results indicate that IV oxycodone maintains greater levels of operant responding when females are in metestrus/diestrus as compared to proestrus/estrus, however this did not coincide with significantly-increased oxycodone intake during metestrus/diestrus under the present testing conditions.

**Figure 3:**
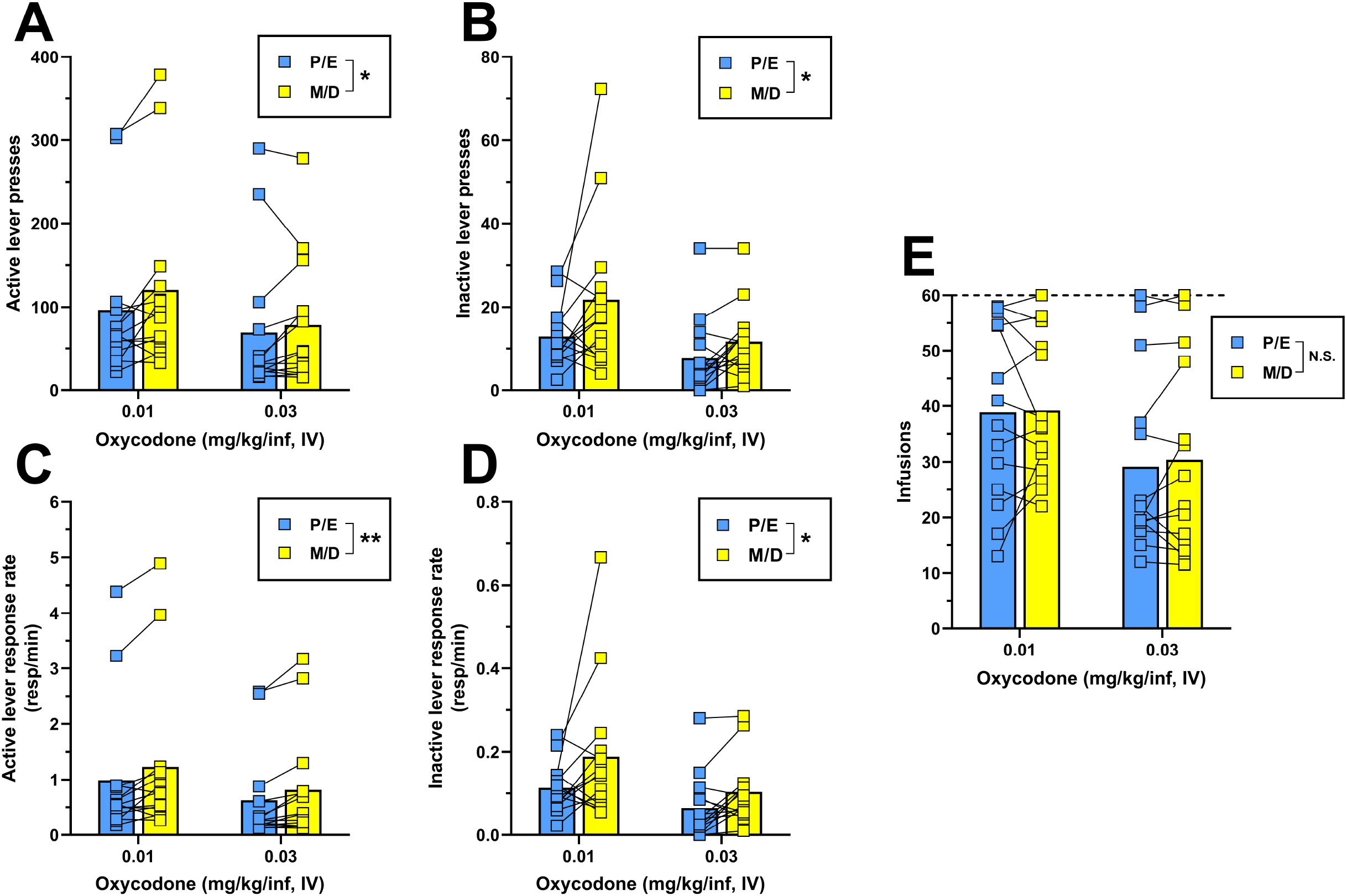
Comparison of IV oxycodone self-administration in female rats during proestrus/estrus vs. metestrus/diestrus. Female (n = 14) Long-Evans rats self-administered 0.03 mg/kg/inf oxycodone for 8 sessions (sessions 1-8) followed by 10 sessions at 0.01 mg/kg/inf (sessions 9-18) under a FR1 schedule of reinforcement. (**A**) Active lever presses, (**B**) inactive lever presses, (**C**) active lever response rate, (**D**) inactive lever response rate, and (**E**) oxycodone infusions earned during proestrus/estrus or metestrus/diestrus, calculated from sessions 5-8 (0.03 mg/kg/inf) and sessions 13-18 (0.01 mg/kg/inf) of oxycodone self-administration (see Fig. 2). The horizontal dashed line in **E** indicates the maximum number of reinforcers allowed. Data are presented as individual data points with connecting lines representing within-subject repeated sampling across estrous cycle phases, superimposed over bars depicting mean values. * p < 0.05, ** p < 0.01, main effect of estrous cycle phase (proestrus/estrus vs. metestrus/diestrus). “P”, proestrus; “E”, estrus; “M”, metestrus; “D”, diestrus; “N.S.”, not significant.

### 3.3 Reinstatement of oxycodone seeking is not modulated by biological sex or estrous cycle

Our final objective was to examine whether there are sex differences and/or estrous cycle-dependent effects on stress-induced or cue-induced reinstatement of oxycodone-seeking behavior. Male and female rats that completed the oxycodone self-administration study described above (8 sessions of 0.03 mg/kg/inf followed by 10 sessions of 0.01 mg/kg/inf) also served as subjects for these reinstatement tests. On the day following the last 0.01 mg/kg/inf oxycodone self-administration session, animals began extinction training until, over two consecutive sessions, the response rate on the active lever was < 30% of their individual mean rate of responding during the last 3 sessions of 0.01 mg/kg/inf oxycodone self-administration. Upon satisfaction of extinction criteria, each animal (male, n = 6; female, n = 13) was first subjected to a footshock-induced reinstatement test, followed by additional extinction training and a cue-induced reinstatement test (see Materials and Methods for additional details). Females were randomly assigned to be tested for footshock- or cue-induced reinstatement while in either a proestrus/estrus or metestrus/diestrus phase of the estrous cycle, with phases identified and targeted as described in the Materials and Methods.

Animals took between 5 – 21 sessions to reach extinction criteria prior to the footshock reinstatement test, with one-way ANOVA indicating no significant difference in the number of extinction sessions required across groups (F_(2,14)_ = 1.56, p = 0.244; data not shown). Estrous cycle phase detection in female subjects on the day of footshock-induced reinstatement testing was as follows: proestrus, n=1/13; estrus, n=6/13; metestrus, n=2/13; diestrus, n=4/13. Surprisingly, exposure to intermittent unpredictable footshock immediately prior to the reinstatement test session did not significantly increase either active or inactive lever presses above extinction levels (**Fig. 4A-B**), as indicated by a lack of main effect of reinstatement phase via two-way ANOVA (active lever, F_(1,16)_ = 0.37, p = 0.550; inactive lever, F_(1,16)_ = 0.91, p = 0.353). The ANOVA also reported no main effect of sex/estrous cycle on active lever responding (F_(2,16)_ = 1.49, p = 0.255) or inactive lever responding (F_(2,16)_ = 1.91, p = 0.180), nor were there significant reinstatement phase × sex/estrous cycle interactions for either lever (active lever, F_(2,16)_ = 1.36, p = 0.286; inactive lever, F_(2,16)_ = 1.34, p = 0.291). Overall, these results indicate that footshock exposure did not elicit appreciable oxycodone-seeking in any of the three experimental groups (**Fig. 4A-B**).

**Figure 4:**
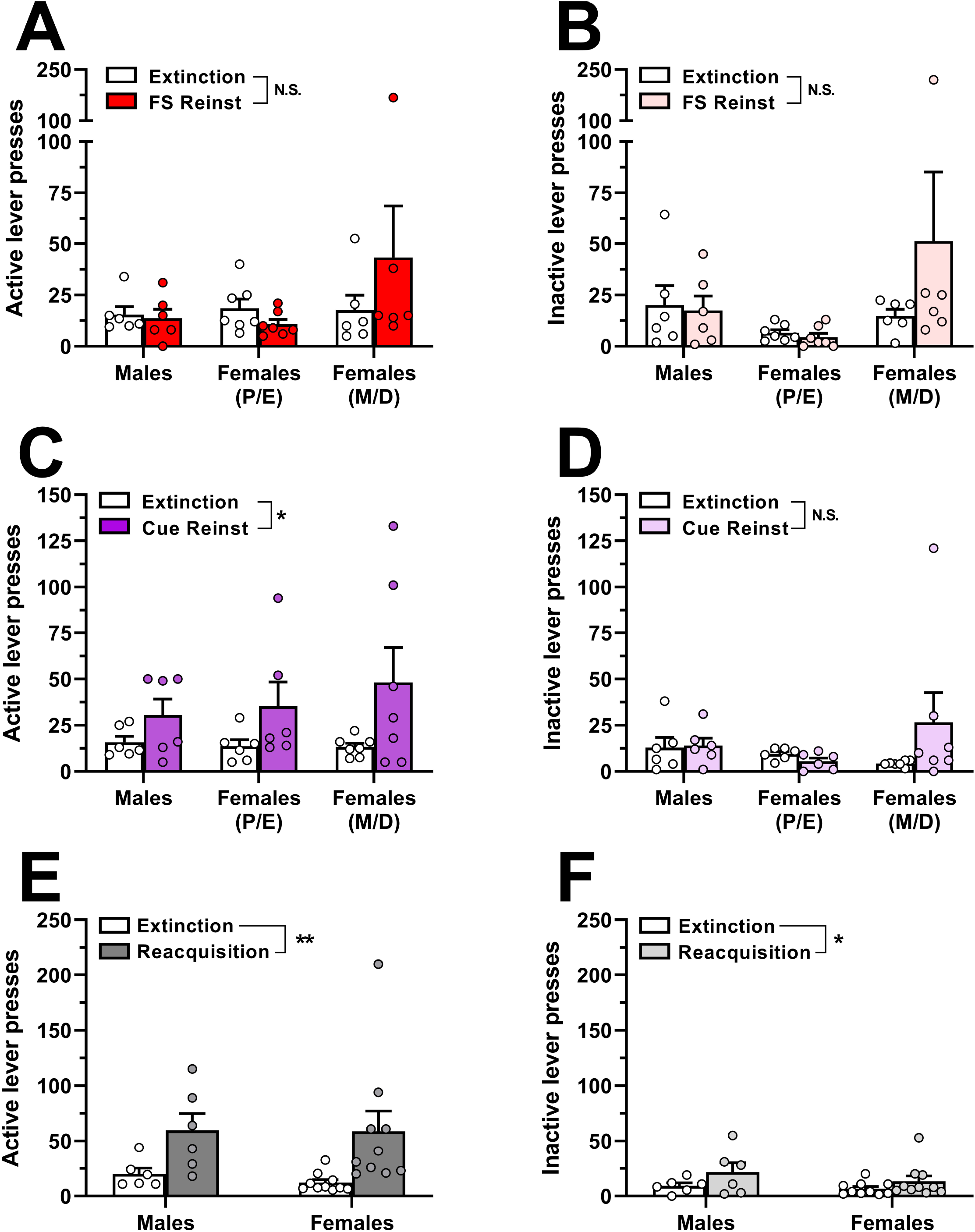
Reinstatement of oxycodone-seeking behavior and reacquisition of oxycodone self-administration in male and female rats. Male and female Long-Evans rats that had previously self-administered oxycodone (see Fig. 2 for details) underwent extinction training followed by stress-induced and cue-induced reinstatement tests. (**A**) Active lever presses and (**B**) inactive lever presses in the last 2 sessions of extinction training (empty bars and symbols) and during a subsequent test session of footshock-induced reinstatement (red/pink bars and symbols). (**C**) Active lever presses and (**D**) inactive lever presses in the last 2 sessions of extinction training (empty bars and symbols) and during a subsequent test session of cue-induced reinstatement (purple/violet bars and symbols). (**E**) Active lever presses and (**F**) inactive lever presses in the last 2 sessions of extinction training (empty bars and symbols) and during a subsequent self-administration session with 0.01 mg/kg/inf oxycodone available (dark gray/light gray bars and symbols). Data are presented as scatter plots with individual subjects represented by circles, superimposed over bars depicting group mean ± SEM values. * p < 0.05, ** p < 0.01, main effect of (**C**) reinstatement phase or (**E, F**) reacquisition. “FS”, footshock; “Reinst”, reinstatement; “P”, proestrus; “E”, estrus; “M”, metestrus; “D”, diestrus; “N.S.”, not significant. n = 6-10/group.

Following the footshock-induced reinstatement test session, each animal was once again subjected to extinction training until extinction criteria were newly satisfied. All rats reached extinction criteria in ≤ 6 sessions. The faster acquisition of extinction here as compared to initial extinction entrainment prior to the footshock reinstatement test is likely due to the animals’ prior experience with extinction training. Nevertheless, each of the experimental groups reached extinction criteria within a similar number of sessions (F_(2,15)_ = 0.12, p = 0.885; data not shown). Estrous cycle phase in female subjects on the day of cue-induced reinstatement testing was as follows: proestrus, n=0/13; estrus, n=6/13; metestrus, n=1/13; diestrus, n=6/13. In contrast to the lack of oxycodone seeking elicited by footshock exposure in the first reinstatement test, reintroduction of response-contingent cues previously paired with oxycodone infusions increased responding selectively on the previously active lever (**Fig. 4C-D**). Two-way ANOVA of active lever presses indicated a significant main effect of reinstatement phase (F_(1,16)_ = 7.77, p = 0.013) but not for sex/estrous cycle (F_(2,16)_ = 0.28, p = 0.761) nor was there a significant reinstatement phase × sex/estrous cycle interaction (F_(2,16)_ = 0.50, p = 0.614). Inactive lever responding did not vary by reinstatement phase (F_(1,16)_ = 1.17, p = 0.295) or sex/estrous cycle (F_(2,16)_ = 0.48, p = 0.625) nor was there a significant reinstatement phase × sex/estrous cycle interaction (F_(2,16)_ = 1.91, p = 0.181). Collectively, these results suggest that reintroduction of IV oxycodone-paired cues elicits oxycodone-seeking behavior, but the magnitude of oxycodone seeking did not differ by biological sex or estrous cycle.

Following the cue-induced reinstatement test, animals that still had patent catheters were re-exposed to daily extinction sessions until criteria were again satisfied and then were allowed to self-administer 0.01 mg/kg/inf oxycodone in a single 2-hr session the next day. All rats extinguished their responding within 7 sessions at rates that did not significantly differ between males and females (t_(14)_ = 1.78, p = 0.10; data not shown). As shown in **Fig. 4E-F**, both male and female rats reliably increased lever-pressing when oxycodone reinforcement was reintroduced. Two-way ANOVA of active lever presses revealed a significant main effect of reacquisition (F_(1,14)_ = 11.88, p = 0.004) but not for sex (F_(1,14)_ = 0.09, p = 0.766) nor was there a significant reacquisition × sex interaction (F_(1,14)_ = 0.08, p = 0.779). Similarly, analysis of inactive lever responding revealed a significant main effect of reacquisition (F_(1,14)_ = 5.64, p = 0.032) but not for sex (F_(1,14)_ = 1.01, p = 0.332) nor was there a significant reacquisition × sex interaction (F_(1,14)_ = 0.49, p = 0.494).

## 4. Discussion

The present study assessed the impact of biological sex and estrous cyclicity on the reinforcing efficacy of IV oxycodone and the reinstatement of previously-extinguished oxycodone-seeking behavior, with three key findings emerging. First, oxycodone functioned as an equally-effective reinforcer in female and male rats under a FR1 schedule of reinforcement. Second, IV oxycodone self-administration maintained higher levels of responding in female rats when they were in the metestrus/diestrus phases of the estrous cycle as compared to proestrus/estrus phases. Finally, exposure to oxycodone-associated cues, but not to acute footshock stress, reinstated lever-pressing in both male and female rats with a prior history of IV oxycodone self-administration, and this behavioral response was not significantly modulated by sex or estrous cycle.

### 4.1 Sex differences in IV oxycodone reinforcement

In the present experiments, IV oxycodone self-administration engendered a typical inverted U-shaped dose-response curve that was centered at 0.01 mg/kg/inf in both male and female rats, consistent with previous reports (Mavrikaki et al., 2017; Mavrikaki et al., 2021). However, despite the fact that females tended to exhibit greater levels of oxycodone-maintained responding in both Experiments 1 and 2, we did not observe sex differences in the reinforcing efficacy of IV oxycodone under the present testing conditions. This was an unexpected result since upward shifts in the dose-response functions for self-administered IV fentanyl and heroin have been demonstrated in female vs. male rats (Cicero et al., 2003; Townsend et al., 2019), and several studies have similarly reported enhanced reinforcing efficacy for IV or oral oxycodone in female rats or mice as compared to males (Mavrikaki et al., 2017; Fulenwider et al., 2020; Kimbrough et al., 2020; Phillips et al., 2020; Zanni et al., 2020). However, others have reported either no sex difference in oxycodone reinforcement, or enhanced reinforcing efficacy in males during food restriction (Mavrikaki et al., 2017; Collins et al., 2020; Mavrikaki et al., 2021). There are numerous procedural differences among these studies that render it challenging to identify specific variables that contribute to these disparate effects. However, it is noteworthy that among studies specifically employing IV oxycodone self-administration procedures, sex differences were reported under conditions of fixed-ratio requirements higher than FR1 (Mavrikaki et al., 2017; Mavrikaki et al., 2021) or using a FR1 schedule during extended access to oxycodone (12 hr/day) (Kimbrough et al., 2020), but not when a short-access (1-2 hr/day) FR1 schedule was utilized to determine a dose-response function (Mavrikaki et al., 2017). It is therefore possible that sex differences in IV oxycodone reinforcement are most readily identifiable under specific testing conditions in which noncontinuous schedules of reinforcement are applied, a phenomenon that has been observed for other drugs of abuse such as cocaine and heroin (Lynch, 2006). Future studies are needed to clarify the experimental parameters under which sex differences in the reinforcing efficacy of IV oxycodone may emerge, and to ascertain whether consequent sex-dependent differences in responding are specific to oxycodone or are generalizable across multiple classes of reinforcing stimuli. We further acknowledge that limiting oxycodone reinforcement to 60 infusions within the short-access 2-hr session may also have impeded the detection of a sex difference specifically on the measure of oxycodone reinforcers earned, because some female subjects earned the maximum number of infusions when 0.003 or 0.01 mg/kg/inf was available, thus imposing an artificial ceiling upon the dose-response curve where infusions earned are the dependent variable on the ordinate. Future studies focused on sex differences in IV oxycodone self-administration should take note of this and implement parameters that might allow for greater separation in responding as well as reinforcers earned across doses and between sexes. For example, one might consider increasing the FR requirement as noted above, extending the duration of drug access, and/or foregoing limitations on the number of allowable infusions per session. It is important to note however that the analysis of response rate, which as an index of reinforcing efficacy is relatively unaffected by reinforcer limitations, did not detect a significant sex difference in the reinforcing effects of IV oxycodone in the present study.

### 4.2 IV oxycodone-maintained responding varies across the estrous cycle

An important finding in the present study is the attenuation of lever pressing maintained by IV oxycodone self-administration during proestrus/estrus in female rats. To the best of our knowledge, this is the first demonstration of estrous cycle-dependent modulation of the abuse-related effects of oxycodone. It should be noted that a prior study examining the role of estrous cycle on oral oxycodone self-administration in female rats did not detect significant differences in operant responding or oxycodone intake across estrous cycle phases (Fulenwider et al., 2020). However, this discrepancy may be due in part to differences in the route of oxycodone administration, the sample sizes tested, and the regularity of the estrous cycle in experimental subjects, among other incongruent variables. Moreover, the differences in IV oxycodone reinforcement that we observed across the estrous cycle are in agreement with other studies demonstrating estrous cycle-dependent fluctuations in the reinforcing effects of remifentanil (Thorpe et al., 2020) and heroin (Lacy et al., 2016; Smith et al., 2021) in intact female rats, suggesting that this may be a phenomenon shared among many, or perhaps all, mu opioid receptor agonists with abuse liability. Proestrus/estrus was associated with a significant attenuation of responses emitted on both the active and inactive operandum during oxycodone self-administration, raising the possibility that nonspecific disruptions in behavioral output may have contributed to observed changes in responding rather than a selective modulation of oxycodone’s efficacy as a behavioral reinforcer per se. Although we did not explicitly test for estrous cycle-dependent effects on general activity levels, we believe that nonspecific reduction of behavioral output can be ruled out as a contributing factor for several reasons. First, others have reported that basal levels of locomotor activity are not significantly impacted by estrous cyclicity in intact female rats (Craft et al., 2006; Scholl et al., 2019; Gaulden et al., 2021). Second, in the present study, levels of active and inactive lever responding during cue-induced reinstatement tests (in the absence of ongoing drug reinforcement) did not significantly differ between females in proestrus/estrus vs. metestrus/diestrus. Finally, operant responding that is reinforced by a nondrug reinforcer (sucrose pellets) under short-access fixed-ratio or progressive-ratio schedules of reinforcement is not different during either proestrus or estrus as compared to metestrus/diestrus in rats (Hecht et al., 1999; Smith et al., 2021). In light of these findings, we interpret reduced output on the inactive lever when oxycodone is self-administered during proestrus/estrus not as an indication of generalized behavioral suppression, but rather as a consequence of the diminished reinforcing efficacy of oxycodone that manifests as attenuated responding directed to both the active and inactive lever, presumably because a small but reliable amount of inactive responding is observed during IV oxycodone self-administration under short-access, fixed-ratio schedules of reinforcement (present study; Mavrikaki et al., 2017; Mavrikaki et al., 2021).

Proestrus cytology was observed less frequently in vaginal lavage samples than estrus cytology, rendering direct comparison between these cycle phases unfeasible due to an insufficient number of proestrus-associated data points. Our experimental design therefore could not differentiate between the relative contributions of proestrus and estrus to the observed reductions in oxycodone-maintained responding since data from these estrous cycle phases were combined into a single dataset for analyses. It is not surprising that the estrus phase was more frequently detected than the proestrus phase because self-administration sessions took place in the early-to-middle portion of the dark period of the light/dark cycle, and proestrus typically coincides with the latter half of the light cycle, transitioning to estrus shortly after onset of the dark cycle in rats (Becker et al., 2005; Lovick and Zangrossi, 2021). Attenuated reinforcing efficacy for two other opioids, heroin and remifentanil, has recently been reported to occur during late proestrus in intact female rats, and not during estrus (Lacy et al., 2016; Thorpe et al., 2020; Smith et al., 2021). However, it remains unknown whether the reinforcing effects of IV oxycodone would be similarly attenuated during late proestrus. Further studies will be necessary to directly compare rates of IV oxycodone self-administration during the proestrus and estrus phases of the estrous cycle to confirm and will require special consideration to the timing of light/dark cycles so as to maximize detection of both proestrus and estrus phases in a within-subjects design.

### 4.3 Impact of sex and estrous cycle on oxycodone-seeking behavior

The lack of a footshock-induced reinstatement response in the present study was surprising since footshock parameters similar to those used here reliably reinstate operant responding previously maintained by opioid reinforcers (for review; Mantsch et al., 2016; Reiner et al., 2019; Nicolas et al., 2022), including oxycodone (Leri and Burns, 2005; Fulenwider et al., 2020). It should be noted that in the same experimental subjects for which footshock was ineffective, we subsequently observed a significant reinstatement effect that was elicited by oxycodone-associated environmental cues, suggesting that the absence of footshock-induced reinstatement was not related to an inability of the animals to exhibit oxycodone-seeking behavior above extinction levels per se, although low sample size should be acknowledged as a limitation of this experiment. The reasons for the lack of footshock-induced reinstatement here remain unclear, but we cannot rule out the possibility that certain aspects of the experimental design may have contributed to this null result. For example, we extinguished oxycodone self-administration in the absence of the oxycodone-paired cue light that functioned as a conditioned reinforcer during self-administration, but some have noted that footshock-induced reinstatement is most effective when response-contingent drug-paired cues are present during both extinction training and the reinstatement test session (Shelton and Beardsley, 2005; Mantsch et al., 2016; Reiner et al., 2019). Additionally, we note that extinction responding was high in some subjects, especially females, which may have precluded a significant reinstatement effect from being detected. For example, mean operant output during the last three extinction sessions was in the range of 30-50 active lever presses in some female subjects, a level of responding that is similar to that observed during footshock-induced reinstatement of oxycodone seeking where extinction baselines are lower (Leri and Burns, 2005; Fulenwider et al., 2020). Setting extinction criteria for all rats to a specific number of active lever presses (e.g., ≤ 15 presses), rather than as a percentage of each individual subject’s self-administration response rate, may have produced lower levels of extinction responding that would be more conducive to the detection of enhanced responding following footshock exposure. Finally, because footshock-induced reinstatement of opioid seeking exhibits an inverted U-shaped intensity-response function (Shaham, 1996), one might speculate that footshock did not elicit reinstatement in the present study because oxycodone-exposed animals were hyperresponsive to the footshock stimulus, perhaps as a consequence of opioid withdrawal-induced hyperalgesia (Laulin et al., 1998; Celerier et al., 2001; Laboureyras et al., 2014). We cannot definitively rule out this possibility since we did not assess whether animals exhibited hyperalgesia at the time of reinstatement testing, however this explanation seems unlikely because we used a footshock exposure protocol that routinely reinstates seeking for opioids and other drugs of abuse (for review; Mantsch et al., 2016; Reiner et al., 2019; Nicolas et al., 2022). More work will be needed to identify the methodological parameters that reliably engender stress-induced reinstatement of IV oxycodone-seeking behavior in male and female rats before the impact of either biological sex or estrous cyclicity on this phenomenon can be assessed.

In contrast to the effect of acute footshock exposure, the reintroduction of response-contingent oxycodone-paired cues following extinction training resulted in significant oxycodone-seeking behavior in both male and female subjects. This finding is consistent with other studies that have reported cue-elicited oxycodone-seeking responses using the extinction-reinstatement model of relapse in rats, although it is noteworthy that only males were used (Leri and Burns, 2005; Neelakantan et al., 2017; Nawarawong et al., 2019). Ours is therefore the first study to show cue-induced oxycodone seeking in female rats, an important but somewhat expected result given prior demonstrations of robust cue-induced reinstatement in female rats where opioids other than IV oxycodone were used as the primary reinforcer (Smethells et al., 2020; Bakhti-Suroosh et al., 2021; Malone et al., 2021; Nett and LaLumiere, 2022; Towers et al., 2022).

Although the highest levels of cue-induced responding were observed in female subjects, the mean magnitude of cue-induced oxycodone seeking in either of the two female experimental groups did not significantly differ from that of males, suggesting that sex did not robustly impact cue-induced oxycodone-seeking behavior in the present study. This interpretation is in agreement with studies that also found no sex differences in the cue-induced reinstatement of responding previously maintained by oral oxycodone (Phillips et al., 2020), IV fentanyl (Bakhti-Suroosh et al., 2021; Malone et al., 2021; Towers et al., 2022), and IV heroin (Smethells et al., 2020; but see Vazquez et al., 2020). Indeed, a recent review on the role of sex in preclinical models of drug seeking concluded that, at present, there is insufficient preclinical evidence to suggest that sex differences exist with regard to cue- or context-induced reinstatement of opioid seeking, a position with which our results are congruent (Nicolas et al., 2022). It is nevertheless intriguing that the highest levels of cue-induced oxycodone seeking occurred in a subset of female rats irrespective of estrous cycle, raising the possibility that other sex-dependent factors may predispose certain female subjects to cue-elicited opioid seeking.

Investigations into the role of estrous cyclicity on cue-induced reinstatement of opioid seeking have only very recently been initiated. Two studies have reported enhanced cue-induced reinstatement of fentanyl seeking during estrus as compared to non-estrus phases (Bakhti-Suroosh et al., 2021; Towers et al., 2022), while another found no effect of estrous cycle on cue-induced fentanyl seeking (Malone et al., 2021). Moreover, incubated heroin seeking observed on day 30 of forced abstinence did not differ between female rats in proestrus/estrus vs. metestrus/diestrus phases (Mayberry et al., 2022). As noted elsewhere, procedural differences may partially explain the discordant effects between some of these studies (Nicolas et al., 2022), but it is clear from these early investigations that more research is warranted on the topic. To the best of our knowledge, the present study is the first to assess whether cue-induced reinstatement of oxycodone seeking varies across the estrous cycle in female rodents. Using a short-access FR1 IV self-administration procedure with a relatively low dose of oxycodone (0.01 mg/kg/inf) as the primary reinforcer available prior to extinction training, we found that the magnitude of cue-induced oxycodone seeking did not vary across the estrous cycle. A noteworthy limitation is that we did not collect data from females during proestrus. Future research should evaluate and directly compare between each distinguishable phase of the estrous cycle the conditioned reinforcing efficacy of oxycodone-paired cues (e.g., using second-order schedules of reinforcement (Di Ciano and Everitt, 2005)), as well as the capacity of oxycodone-associated cues to reinstate previously-extinguished oxycodone-maintained responding or elicit oxycodone seeking in other models of relapse.

## 5. Conclusions

In summary, we report that IV oxycodone functions as an equally-effective reinforcer in male and female rats under a short-access continuous schedule of reinforcement and, for the first time, demonstrate that oxycodone-maintained responding is highest in females during metestrus/diestrus phases of the estrous cycle and is therefore by comparison attenuated during proestrus/estrus. At present, the mechanisms underlying the estrous cycle-dependent modulations of oxycodone reinforcement remain unknown, although evidence is accumulating to suggest that fluctuations in ovarian hormones, particularly estradiol, may be responsible for the latter (Smith et al., 2021; Smith et al., 2022). There is a clear need for future studies to examine these potential mechanistic underpinnings, especially in light of the enhanced prevalence of prescription opioid use and misuse in women as compared to men (Unick et al., 2013; Han et al., 2017; Serdarevic et al., 2017). By contrast, using an extinction/reinstatement model of drug relapse, we found no evidence that cue-induced reinstatement of oxycodone seeking differs between male and female rats, nor does it vary between estrus vs. metestrus/diestrus phases of the estrous cycle. Collectively, our results suggest that ovarian hormone levels should be taken into account when studying the primary reinforcing effects of oxycodone in both preclinical and clinical settings.

## AUTHOR CONTRIBUTIONS

DM and IW contributed to initial conception and design of the study. NH, IW, CG, CC, and DM performed experiments and collected data. DM performed the statistical analyses. NH, IW, and DM contributed to the first draft of the manuscript. DM edited and finalized the manuscript. NH, IW, CG, CC, and DM all read and approved the final manuscript prior to submission.

## ACKNOWLEDGEMENTS

The authors would like to thank the NIDA Drug Supply Program for generously providing oxycodone hydrochloride.

