## Supplementary Material for "Effects of sex and estrous cycle on intravenous oxycodone self-administration and the reinstatement of oxycodone-seeking behavior in rats"

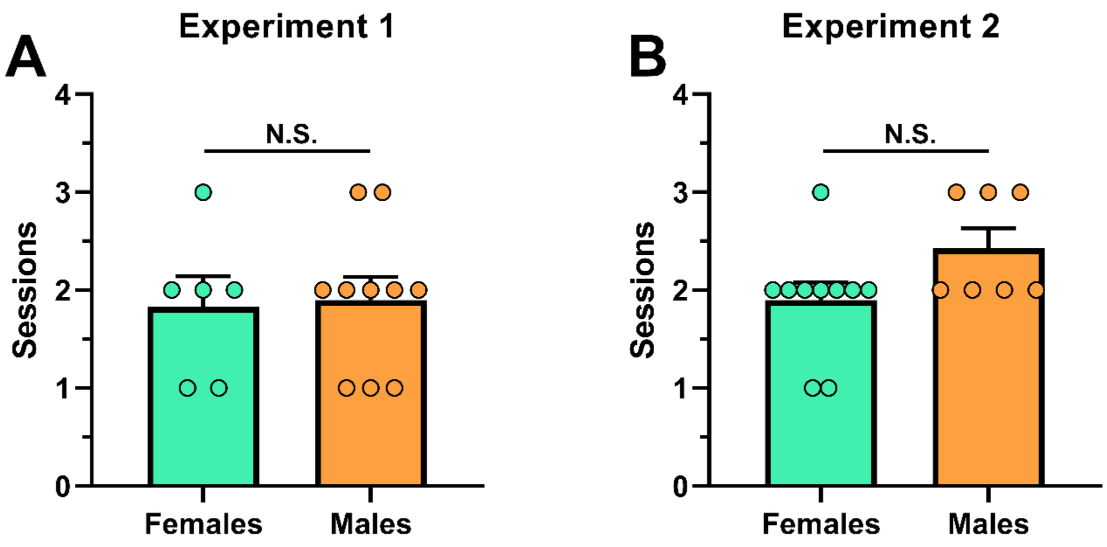


**Supplemental Figure 1: Rates of acquisition for lever-press training under a FR1 schedule of food-maintained responding.** Male and female Long-Evans rats used as subjects in Experiment 1 (**A**) or Experiment 2 (**B**) were first exposed to food presentation according to a FR1 schedule of reinforcement (see Materials and Methods for additional details). Shown are the number of sessions required to satisfy training criteria in male and female subjects that successfully completed food training. Data are presented as scatter plots with individual subjects represented by circles, superimposed over bars depicting group mean ± SEM values. “N.S.”, not significant.

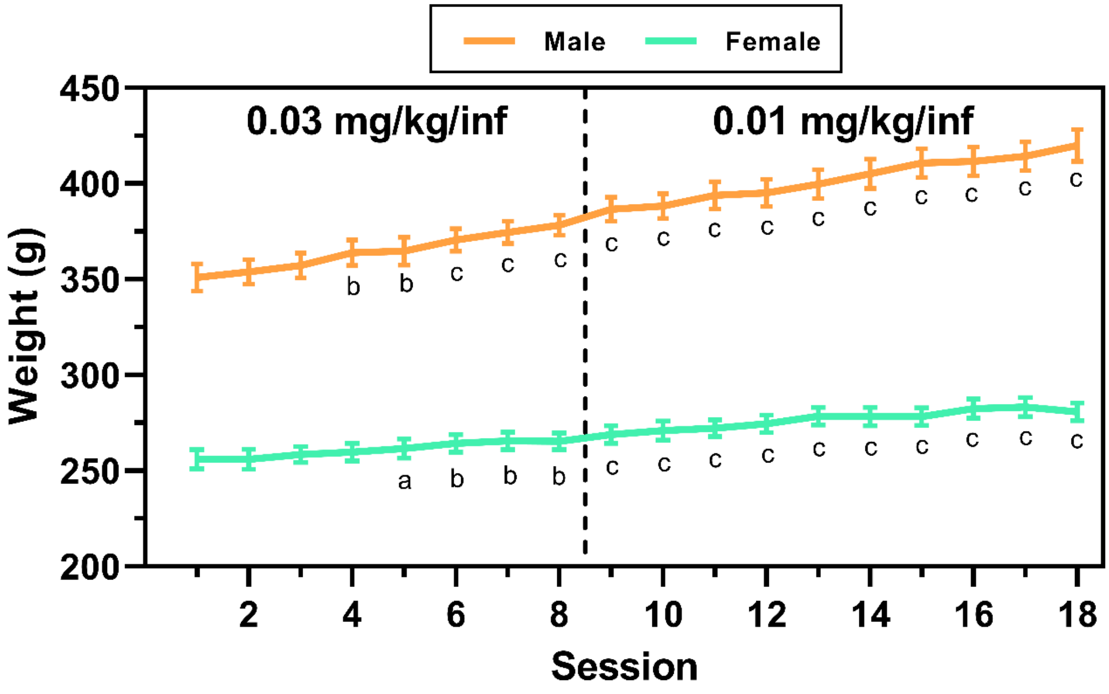


**Supplemental Figure 2: Daily body mass during IV oxycodone self-administration in male and female rats used in Experiment 2.** Male (n = 8) and female (n=14) Long-Evans rats used as subjects in Experiment 2 were weighed immediately prior to each oxycodone self-administration session. The vertical dashed line indicates the transition from 0.03 to 0.01 mg/kg/inf oxycodone availability. Data are presented as mean ± SEM weight in grams. “a” $p < 0.01$, “b” $p < 0.001$, “c” $p < 0.0001$, Dunnett’s test compared to session 1 within sex.
